# Learning spatiotemporal signals using a recurrent spiking network that discretizes time

**DOI:** 10.1101/693861

**Authors:** Amadeus Maes, Mauricio Barahona, Claudia Clopath

## Abstract

Learning to produce spatiotemporal sequences is a common task the brain has to solve. The same neural substrate may be used by the brain to produce different sequential behaviours. The way the brain learns and encodes such tasks remains unknown as current computational models do not typically use realistic biologically-plausible learning. Here, we propose a model where a spiking recurrent network of excitatory and inhibitory biophysical neurons drives a read-out layer: the dynamics of the recurrent network is constrained to encode time while the read-out neurons encode space. Space is then linked with time through plastic synapses that follow common Hebbian learning rules. We demonstrate that the model is able to learn spatiotemporal dynamics on a timescale that is behaviourally relevant. Learned sequences are robustly replayed during a regime of spontaneous activity.

**Author summary:** The brain has the ability to learn flexible behaviours on a wide range of time scales. Previous studies have successfully build spiking network models that learn a variety of computational tasks. However, often the learning involved is not local. Here, we investigate a model using biological-plausible plasticity rules for a specific computational task: spatiotemporal sequence learning. The architecture separates time and space into two different parts and this allows learning to bind space to time. Importantly, the time component is encoded into a recurrent network which exhibits sequential dynamics on a behavioural time scale. This network is then used as an engine to drive spatial read-out neurons. We demonstrate that the model can learn complicated spatiotemporal spiking dynamics, such as the song of a bird, and replay the song robustly.

## Introduction

Neuronal networks perform flexible computations on a wide range of time scales. While individual neurons operate on the millisecond time scale, behaviour typically spans from a few milliseconds to hundreds of milliseconds and longer. Building functional models that bridge this time gap is of increasing interest [Abbott et al., 2016], especially now that the activity of many neurons can be recorded simultaneously [Jun et al., 2017, Maass, 2016]. Many tasks and behaviours essentially have to produce spatiotemporal sequences. For example, songbirds produce their songs through a specialized circuit. Neurons in the HVC nucleus burst sparsely at very precise times to drive the robust nucleus of the arcopallium which in its turn drives motor neurons [Hahnloser et al., 2002, Leonardo and Fee, 2005]. In addition, sequential neuronal activity is recorded in various brain regions for different motor tasks [Pastalkova et al., 2008, Itskov et al., 2011, Harvey et al., 2012, Peters et al., 2014, Adler et al., 2019]. While the different tasks possibly involve different sets of muscles, the underlying computation on a more fundamental level might be similar [Rhodes et al., 2004].

Theoretical and computational studies have shown that synaptic weights of recurrent networks can be set appropriately such that dynamics on a wide range of time scales is produced [Litwin-Kumar and Doiron, 2014, Zenke et al., 2015, Tully et al., 2016]. In general, these synaptic weights are engineered to generate a range of interesting dynamics: slow-switching dynamics [Schaub et al., 2015] and different types of sequential dynamics [Chenkov et al., 2017, Billeh and Schaub, 2018, Setareh et al., 2018]. However, it is unclear how the brain learns these dynamics as most of the current approaches use non biologically-plausible ways to set or “train” the synaptic weights. For example, FORCE training [Sussillo and Abbott, 2009, Laje and Buonomano, 2013, Nicola and Clopath, 2017] or backpropagation through time use non-local information either in space or in time.

Here, we propose to learn a task in a spiking recurrent network which drives a read-out layer. All synapses are plastic under typical Hebbian learning rules [Clopath et al., 2010, Litwin-Kumar and Doiron, 2014]. We train the recurrent network to generate a sequential activity which serves as a temporal backbone. This sequential activity is generated by clusters of neurons activated one after the other. As clusters are highly recurrently connected, each cluster undergoes reverberating activity that lasts longer than neural time scale. Therefore, the sequential activation of the clusters leads to a sequence long enough to be behaviourally relevant. This allows us to bridge the neural and the behavioural time scales. We then use Hebbian learning to learn the target spatiotemporal dynamics in the read-out neurons. Thus, the recurrent network encodes time and the read-out neurons encode ‘space’. ‘Space’ can mean spatial position, but it can also mean frequency or a more abstract state space. Similar to the liquid state-machine, we can learn different dynamics in parallel in different read-out populations [Jaeger et al., 2007]. We show that learning in the recurrent network is stable during spontaneous activity and that the model is robust to synaptic failure.

## Results

In the following we will build our model step-by-step in a bottom-up manner.

### Model architecture

The model consists of two separate parts (Fig 1A). The first part is a recurrent network which serves to encode time in a discretized manner. The second is a read-out layer. The read-out layer learns a multivariate signal of time, *t* → *ϕ*(*t*). The neurons in the read-out layer encode the *D* different dimensions of the signal, *ϕ*(*t*) = [*ϕ*_1_(*t*), *ϕ*_2_(*t*), *…, ϕ*_*D*_(*t*)]. Time is discretized in the recurrent spiking network by *C* clusters of excitatory neurons, *t* = [*t*_0_, *t*_1_, …, *t*_*C*_]. This means that at each time only a subset of neurons spike, i.d. the neurons belonging to the same cluster. Given that the first cluster is active during time interval [*t*_0_, *t*_1_], cluster *i* will be active during time interval [*t*_*i-*1_, *t*_*i*_]. These excitatory clusters drive the read-out neurons through all-to-all feedforward connections. At time *t* ∈ [*t*_*i-*1_, *t*_*i*_] the read-out neurons are solely driven by the neurons in cluster *i*, with some additional background noise. This discretization enables Hebbian plasticity to bind the read-out neurons to the neurons active in the relevant time-bin. The read-out neurons are not interconnected and receive input during learning from a set of excitatory supervisor neurons. In previous models, the learning schemes are typically not biologically plausible because the plasticity depends on non-local information. Here however, we use the voltage-based STDP plasticity rule at the excitatory to excitatory neurons (Fig 1B). This is paired with weight normalization in the recurrent network (Fig 1C) and weight dependent potentiation in the read-out synapses (Fig 1D). Additionally, inhibitory plasticity [Vogels et al., 2011] prevents runaway dynamics.

**Figure 1:**
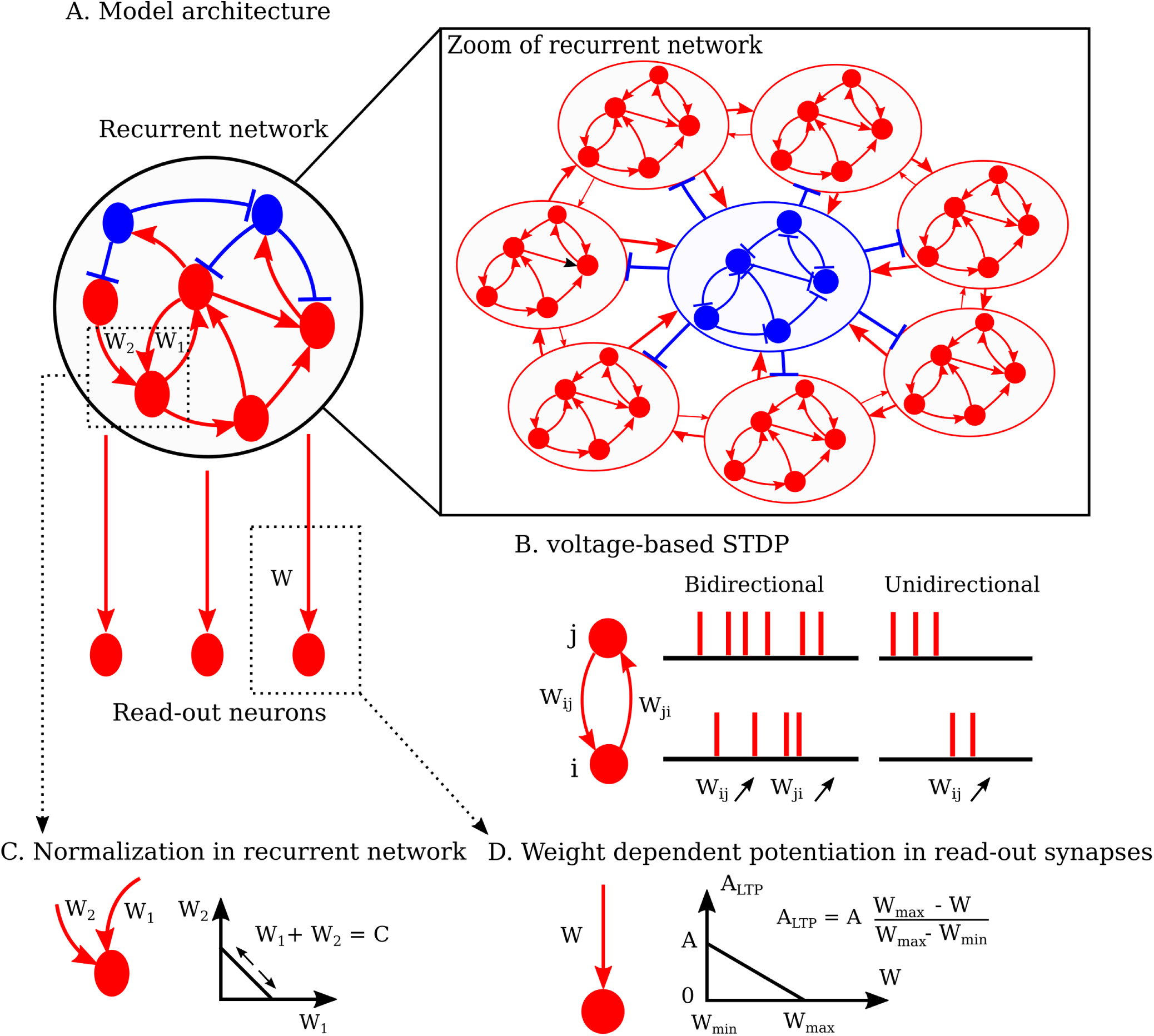
(A.) The recurrent network consists of both inhibitory (in blue) and excitatory (in red) neurons. The connectivity is sparse in the recurrent network. The temporal backbone is established in the recurrent network after a learning phase. Inset: zoom of recurrent network showing the macroscopic recurrent structure after learning, here for 7 clusters. The excitatory neurons in the recurrent network project all-to-all to the read-out neurons. The read-out neurons are not interconnected. (B.) All excitatory to excitatory connections are plastic under the voltage-based STDP rule (see Methods for details). The red lines are spikes of neuron *j* (top) and neuron *i* (bottom). When neurons *j* and *i* are very active together, they form bidirectional connections strengthening both *W*_*ij*_ and *W*_*ji*_. Connections *W*_*ij*_ are unidirectionally strengthened when neuron *j* fires before neuron *i*. (C.) The incoming excitatory weights are *L*1 normalized in the recurrent network, i.d. the sum of all incoming excitatory weights is kept constant. (D.) Potentiation of the plastic read-out synapses is linearly dependent on the weight. This gives weights a soft upper bound.

### A recurrent network that encodes discrete time

Learning happens in a two-stage process. In the first stage, a feedforward weight structure is embedded into the recurrent network. The excitatory neurons are divided into disjoint clusters and neurons in the same cluster share the same input. The recurrent network is initialized as a balanced network with random connectivity. To embed a feedforward structure, the network is stimulated in a sequential manner (Fig 2A). Neurons in cluster *i* each receive external Poisson spike trains (rate of 18 k*Hz* for 10 m*s*). After this, there is a time gap where no clusters receive input (5 m*s*). This is followed by a stimulation of cluster *i*+1. This continues until the last cluster is reached and then it links back to the first cluster (i.e. a circular boundary condition). During the stimulation, neurons in the same cluster fire spikes together, which will strenghten the intra-cluster connections bidirectionally through the voltage-based STDP rule [Clopath et al., 2010, Ko et al., 2013]. Additionally, there is a pre/post pairing between adjacent clusters. The weights from cluster *i* to cluster *i* + 1 strengthen unidirectionally (Fig 2B). When the time gap between sequential stimulations is increased during the training phase, there is no pre/post pairing between clusters anymore. This leads to slow-switching dynamics as opposed to sequential dynamics [Litwin-Kumar and Doiron, 2014, Schaub et al., 2015] (Fig S1). After the weights have converged, the external sequential input is shut-down and spontaneous dynamics is simulated. External excitatory Poisson spike trains without spatial or temporal structure drive the spontaneous dynamics. During that time, the sequence of the clusters reactivates spontaneously and ensures that both the intra-cluster and the feedforward connections remain strong. The connectivity pattern is therefore stable (Fig S2). The feedforward weight embedding changes the spectrum of the recurrent weight matrix. The dominant eigenvalues lay in a circle in the complex plane (Fig 2C). Analysis of a simplified linear rate model shows that the imaginary parts linearly depend on the strength of the feedforward embedding (see supplementary material, Fig S3). Intuitively, this means that the imaginary parts of the dominant eigenvalues determine the period of the sequential activity (Fig S2). Under spontaneous activity, each cluster is active for about 14 m*s*, which is due to the recurrent connectivity within the cluster. A large adaptation current counteracts the recurrent reverberating activity to turn the activity reliably off. Therefore, as each cluster is spontaneously active in a sequence, that leads to a sequence length that reaches behavioural time scales (Fig 2D). In summary, the feedforward block connectivity structure embedded in the recurrent network results in a sequential dynamics that discretizes time.

**Figure 2:**
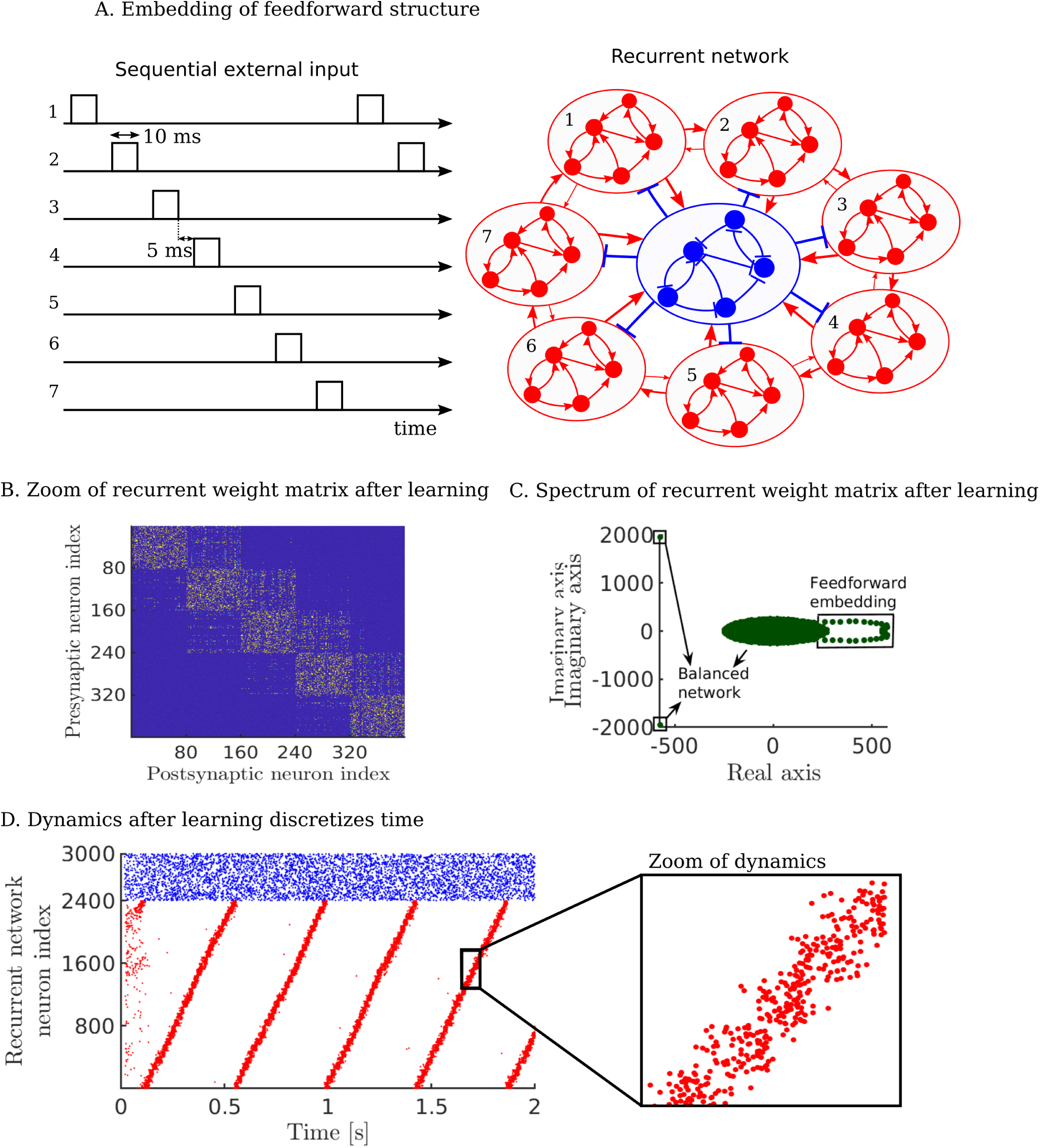
Learning a sequential dynamics that is stable under plasticity. (A.) The excitatory neurons receive sequential clustered inputs. (B.) The weight matrix after training shows the feedforward embedding on the upper diagonal (only the first five clusters displayed), e.g. neurons in cluster 1 are highly bidirectionally connected and also project to neurons in cluster 2, same for cluster 2 to cluster 3 etc. (C.) The spectrum of the full weight matrix after training. The dominant eigenvalues lay in a circle in the complex plane indicating the feedforward embedding on top of the diagonal clustering. (D.) Raster plot of the total network consisting of 2400 excitatory (in red) and 600 inhibitory (in blue) neurons. The spontaneous dynamics exhibits a stable periodic trajectory. The excitatory clusters discretize time and the network has a period of about 420 m*s*. Inset: zoom of the raster plot.

### Learning a non-markovian sequence

After the temporal backbone in the recurrent network is established, we now learn a simple yet non-markovian sequence in the read-out neurons. During training, the read-out neurons receive additional input from supervisor neurons and from interneurons (Fig 3A). The supervisor neurons receive external Poisson input. The rate of the Poisson input is modulated by the target sequence that needs to be learned (Fig 3B). The target sequence is composed of different states (*A, B* or *C*) that are activated in the following order *ABCBA*. This is a non-markovian state sequence because the transition from state *B* requires knowledge about the previous state. Each neuron in the supervisor is encoding one state. This task is non-trivial and it is unclear how to solve this problem in a generic recurrent spiking network with local plasticity rules. However, separating time and space solves this in a natural way. Our recurrent network (Fig 2) is used here, where the first cluster is activated at time *t*. At the same time *t*, the supervisor is also activated. The underlying assumption is that a starting signal activates both the first cluster of the recurrent network and the external input to the supervisor neurons at the same time. The read-out neurons are activated by the supervisor neurons. Due to the voltage-based plasticity, synapses are potentiated from neurons in the recurrent network and read-out neurons that fire at the same time. After the training period, the interneurons and supervisor neurons stop firing (Fig 3C). The target sequence is now stored in the read-out weight matrix (Fig 3D). As clusters in the recurrent network spontaneously reactivate during spontaneous activity in a sequential manner, they also reactivate the learned sequence in the read-out neurons. The spike sequence of the read-out neurons is a noisy version of the target signal (Fig 3E). Learning the same target signal several times results in slightly different read-out spike sequences each time (Fig S4). The firing rates of neurons in the read-out corresponds to the target sequence (Fig 3F). In summary, our results show that the model is able to learn simple but non-trivial spike sequences.

**Figure 3:**
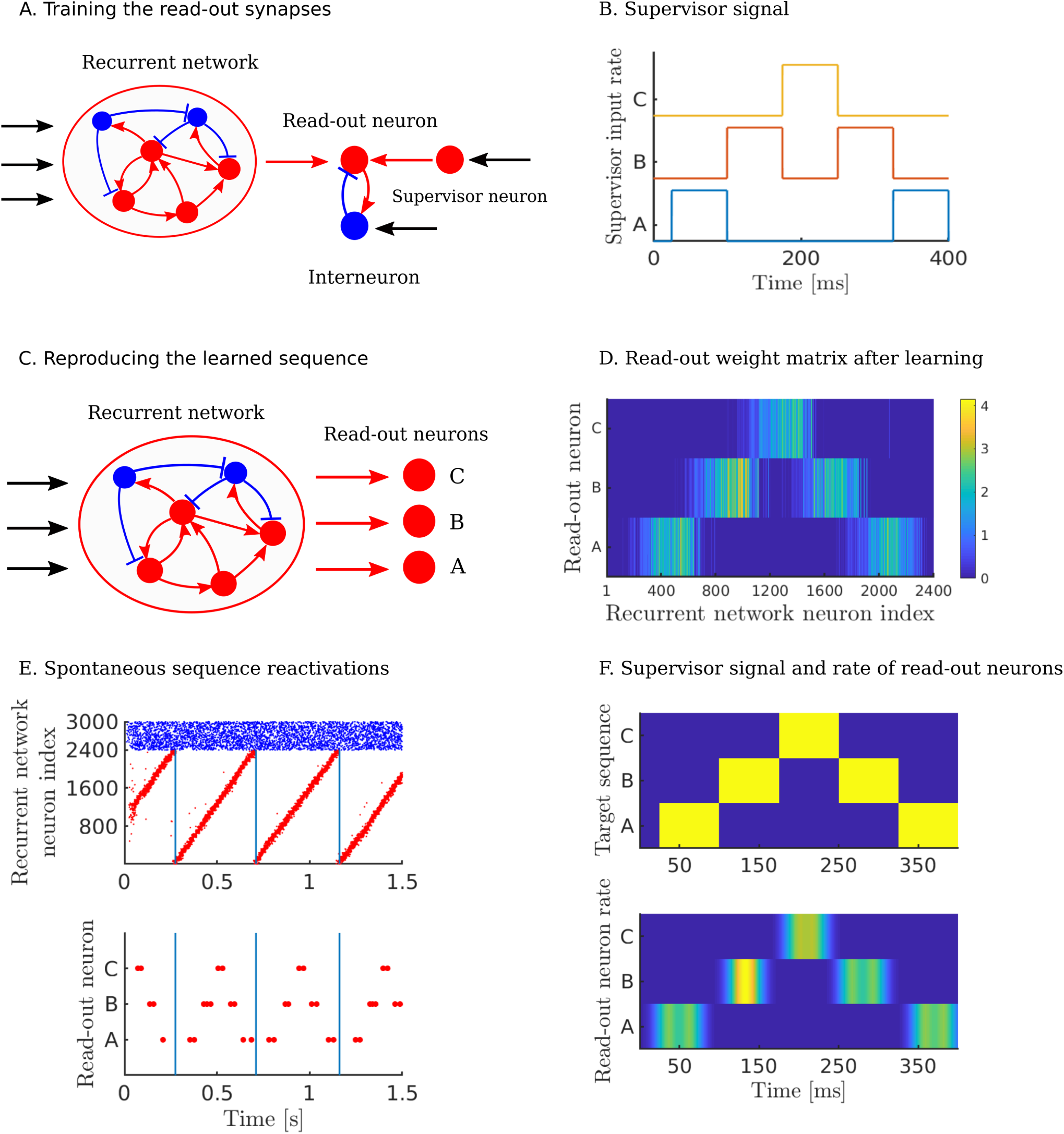
Learning a non-Markovian sequence. (A.) Excitatory neurons in the recurrent network are all-to-all connected to the read-out neurons. The read-out neurons receive additional excitatory input from the supervisor neurons and inhibitory input from interneurons. The supervisor neurons receive spike trains that are drawn from a Poisson process with a rate determined by the target sequence. The read-out synapses are plastic under the voltage-based STDP rule. (B.) The rate of the input signal to the supervisor neurons *A, B* and *C*. The supervisor sequence is *ABCBA* where each letter represents a 75 m*s* external stimulation of 10 k*Hz* of the respective supervisor neuron. (C.) After 12 seconds of learning, the supervisor input and plasticity are turned off. The read-out neurons are now solely driven by the recurrent network. (D.) The read-out weight matrix *W*^*RE*^ after learning. (E.) Under spontaneous activity, the spikes of recurrent network (top) and read-out (bottom) neurons. Excitatory neurons in the recurrent network reliably drive sequence replays. (F.) The target rate (above) and the rate of the read-out neurons (below) computed using a one sequence replay and normalized to [0, 1]. The spikes of the read-out neurons are convolved with a Gaussian kernel with a width of ∼ 12 m*s*.

### Properties of the model

We then investigate the robustness of our model by silencing neurons in the recurrent network after learning. The first type of robustness we test is a robustness of the read-out. To study this, the presynaptic spike trains from the recurrent network are slightly perturbed. We compared two different networks. 1) A large network (where 2400 excitatory neurons are grouped in 30 clusters) and 2) a small network (where 600 excitatory neurons are grouped in 30 clusters). In order to have flexibility, we did not simulate our recurrent network dynamics but simply forced neurons in cluster *i* to spike randomly in time bin *i*. As before, we learn the simple *ABCBA* sequence in the readout neurons (Fig 4A). Note that the larger network learns quicker than the smaller one. The strengths of the read-out synapses of the smaller clusters need to be larger to be able to drive the read-out neurons. After learning, 4 random neurons are silenced in each cluster at two different time points. While the effect of silencing these neurons is very small in the larger network, it is very visible in the smaller network. Multiple spikes disappear at each read-out neuron. These results show that, not surprisingly, larger clusters drive read-out neurons more robustly.

**Figure 4:**
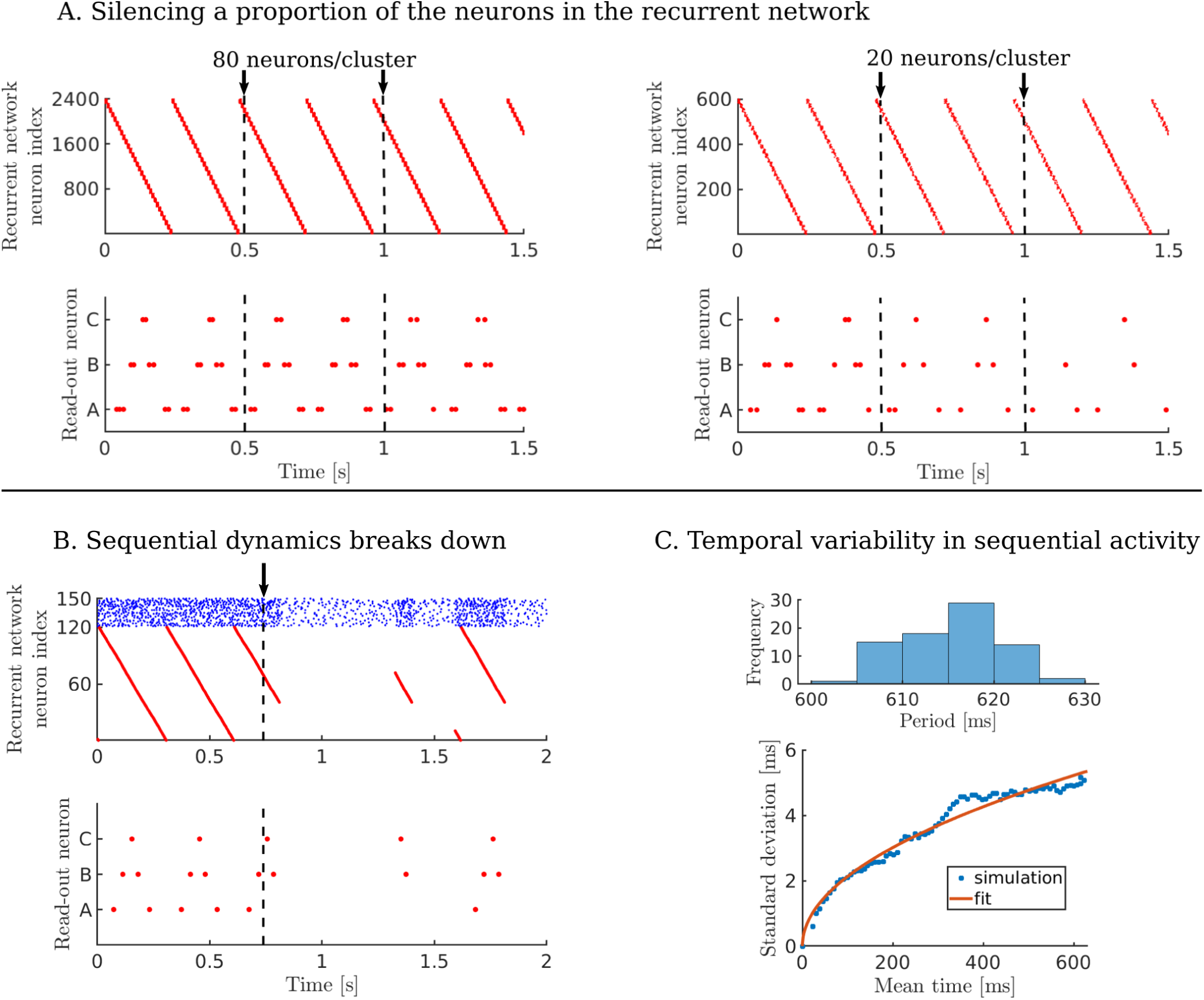
Robustness and time variability of the model. (A.) The sequential dynamics of the large (2400 neurons, left) and small (600 neurons, right) network are artificially elicited. In the large network, 2400 neurons are split in 30 clusters of 80 neurons, as before. In the small network, the 600 neurons are split in 30 clusters of 20 neurons. The neurons of cluster *i* fire in time bin *i*, each time bin has a fixed duration of 8 m*s*. After learning the non-markovian sequence *ABCBA* where each letter indicates a 40 m*s* stimulation, neurons are silenced. The large network learns for 15 seconds and the small network learns for 60 seconds. At *t* = 500 m*s* and at *t* = 1000 m*s*, 4 neurons are randomly silenced in each cluster. Top: spike raster of the excitatory neurons of the recurrent network. Bottom: spike raster of the read-out neurons. (B.) 120 excitory neurons are connected in one large chain and driven by a low external input 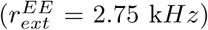. After 200 seconds of learning, spontaneous dynamics is simulated. At *t* = 750 m*s*, a single synapse (the synapse connecting neuron number 41 to neuron number 40) is deleted. The dynamics breaks down and parts of the sequence are randomly activated by the external input. The learned read-out dynamics is fractured in subsequences as a consequence. Top: spike raster of the excitatory neurons of the recurrent network. Bottom: spike raster of the read-out neurons. The maximum synaptic read-out strength is increased, 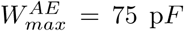, to be able to learn large read-out weights. (C.) Temporal variability in the sequential activity of the recurrent network is measured over 79 trials. A histogram of the periods of the 79 sequences (top) shows an average period of about 615 m*s* and a standard deviation of around 5 m*s*. The standard deviation increases roughly with the square root of time (bottom). The least squares fit for the function 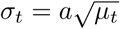 has a root mean squared error of 0.223 m*s. µ*_*t*_ is the mean time that a cluster is activated, *σ*_*t*_ is the standard deviation of the cluster activations and *a* the coefficient of the fit, *a* = 0.213. A linear fit increases the root mean squared error to 0.824 m*s*. The lowest time constant in the plasticity of the read-out synapses is 5 m*s*.

We then wanted to test whether time discretization is important in our model. To that end, we trained our recurrent network with clusters as small as one neuron. In that extreme case, the clusters and recurrent connections effectively disappear and our network becomes a synfire chain with a single neuron in every layer [**?**]. Randomly removing a synapse in the network will break the sequential dynamics (Fig 4B). Although a spatiotemporal signal can be learned in the read-out neurons, the signal is not stable under a perturbation as the synfire-chain is not stable (Fig 4B). In summary, the choice of cluster size is a trade-off between network size on the one hand and robustness on the other hand. Large clusters require a large network but learn faster, are less prone to a failure of the sequential dynamics and drive the read-out neurons more robustly.

The sequential activity in the recurrent network shows some variability in terms of sequence duration. The neural activity does not move along the periodic trajectories with exactly the same speed in each reactivation. We wondered how the variance in our network compared to Weber’s law. According to Weber’s law, the standard deviation of reactions in a timing task grows linearly with time [Gibbon, 1977, Hardy and Buonomano, 2018]. Since our model operates on a time scale that is behaviourally relevant, it is interesting to look at how the variability increases with increasing time. Time is discretized by clusters of neurons that activate each other sequentially, as such the variability increases naturally over time. As in a standard diffusion process, this increase is expected to grow with 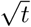 rather than linearly. This is indeed what our network displays (Fig 4C). Here, we increased the period of the recurrent network by increasing the network size (80 excitatory clusters of 80 neurons each, see Methods for details). By doing so, we can look at how the standard deviation of cluster activations grows with time.

### Learning a complex sequence

In the non-markovian target sequence *ABCBA*, the states have the same duration and the same amplitude (Fig 3B). We wanted to test whether we could learn some more complicated sequences. In this second task, the model is trained using a spatiotemporal signal that has components of varying durations and amplitudes. As an example, we use a 600 m*s* meadowlark song (Fig 5A). The spectrogram of the sound is normalized and used as the time- and amplitude-varying rate of the external Poisson input to the supervisor neurons. Each read-out and supervisor neuron encodes a different frequency range. The song is longer than the period of the original recurrent network (Fig 2). Therefore, we trained on the larger network of 6400 excitatory neurons (Fig 4C). After training, the read-out weight matrix reflects the structure of the target sequence (Fig 5B). Under spontaneous activity, the supervisor neurons and interneurons stop firing and the recurrent network drives song replays (Fig 5C). The learned spatiotemporal signal broadly follows the target sequence (Fig 5A). The model performs worse when the target dynamics has fast time-variations. This is because the time-variations are faster than or at the same order as the time discretization in the recurrent network. Thus, we conclude that the model can learn interesting spiking dynamics. The performance is qualitatively better when the changes in the target dynamics are slower than the time discretization in the recurrent network.

**Figure 5:**
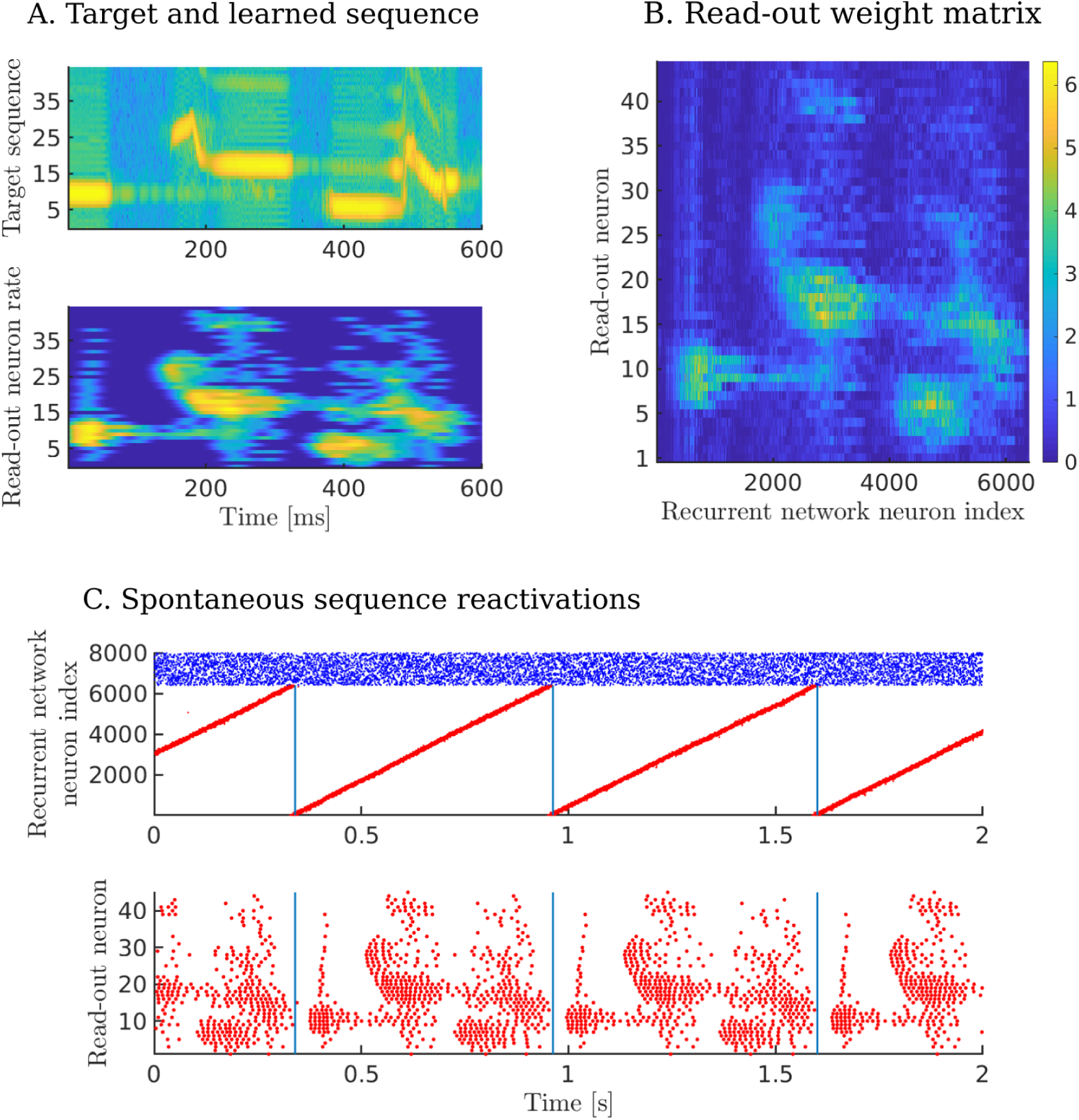
(A.) Target sequence (top). The amplitude shows the rate of the Poisson input to the supervisor neurons and is normalized between 0 and 10 k*Hz*. Rate of read-out neurons for one sample reactivation after learning 6 seconds (bottom). 45 read-out neurons encode the different frequencies in the song. Neuron *i* encodes a frequency interval of [684 + 171*i*, 855 + 171*i*]*Hz*. (B.) The read-out weight matrix after learning 6 seconds. (C.) Sequence replays showing the spike trains of both the recurrent network neurons (top, excitatory neurons in red and inhibitory neurons in blue), and the read-out neurons (bottom).

## Discussion

We proposed an architecture that consists of two parts. The first part is a temporal backbone, implemented by a recurrent network. Excitatory neurons in the recurrent network are organized into disjoint clusters. These clusters are sequentially active by embedding a feedforward connectivity structure into the weights of the network. We showed that ongoing spontaneous activity does not degrade this structure. The set of plasticity rules and sequential dynamics reinforce each other. This was shown for slow-switching dynamics before [Litwin-Kumar and Doiron, 2014] (Fig S1). The stable sequential dynamics provides a downstream linear decoder with the possibility to read time out at the behavioural timescale.

The second part is a set of read-out neurons that encode multiple dimensions of a signal, which we also call “space”. Connecting the first to the second part binds space to time. The read-out neurons learn spike sequences in a supervised manner. More specifically, we investigated a simple state transition sequence and a sequence that has a more complex dynamics. The supervisor sequence is encoded into the read-out weight matrix.

The presented model brings elements from different studies together. Recurrent network models that learn sequential dynamics are widely studied [Fiete et al., 2010, Rajan et al., 2016, Hardy and Buonomano, 2018]. Separate from this, models exist that aim to learn spatiotemporal dynamics [Brea et al., 2013, Nicola and Clopath, 2017, Gilra and Gerstner, 2017]. Combining the two, the dynamics of the recurrent network is exploited by the read-out neurons to perform the computational task. In this paper, we use local Hebbian learning such as the voltage-based STDP [Clopath et al., 2010].

Importantly, the dynamics of the recurrent network spans three time scales. First, individual neurons fire at the fastest milliseconds time scale. Second, one level slower, clusters of neurons fire at a time scale that determines the time discretisation of our temporal backbone, or if you will, the “tick of the clock”, *τ*_*c*_. Third, the slowest time scale is on the level of the entire network, i.e. the period of the sequential activity, *τ*_*p*_. The time scales *τ*_*c*_ and *τ*_*p*_ are dependent on the cluster and network size, the average connection strengths within the clusters and adaptation. Smaller cluster sizes lead to a smaller *τ*_*c*_ and *τ*_*p*_ increases with network size when the cluster size is fixed.

The recurrent network is the “engine” that, once established, drives read-out dynamics. Our model can learn different read-out synapses in parallel and is robust to synapse failure. The robustness is a consequence of the clustered organization of the recurrent network. The clusters in the recurrent network provide a large drive for the read-out neurons while keeping the individual synaptic strengths reasonably small. Smaller clusters on the other hand require larger read-out synaptic strengths and the dynamics are as such more prone to small changes. In the limit, every cluster has exactly one neuron. The sequential dynamics is especially fragile in this case. Learning is faster with more neurons per cluster. Relatively small changes in the synapses are sufficient to learn the target. This is consistent with the intuitive idea that some redundancy in the network can lead to an increased learning speed [Raman et al., 2019].

Previous studies have discussed whether sequences are learned and executed serially or hierarchically [Lashley, 1951]. Our recurrent network has a serial organization. When the sequential activity breaks down halfway, the remaining clusters are not activated anymore. A hierarchical structure would avoid such complete break-downs at the cost of more complicated structure to control the system. Sequences that are chunked in sub-sequences can be learned separately and chained together. When there are errors in early sub-sequences this will less likely affect the later sub-sequences. Evidence for hierarchical structures is found throughout the literature [Sakai et al., 2003, Glaze and Troyer, 2006, Okubo et al., 2015]. The basal ganglia is for example thought to play an important role in shaping and controlling action sequences [Tanji, 2001, Jin and Costa, 2015, Geddes et al., 2018]. Another reason why a hierarchical organization seems beneficial is inherent to the sequential dynamics. The time-variability of the sequential activity grows by approximately 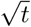. While on a time scale of a few hundreds of milliseconds, this does not yet pose a problem, for longer target sequences this variability would reach and exceed the plasticity time constants. The presented model can serve as an elementary building block of a more complex hierarchy.

In summary, we have shown that a clustered network organization can be a powerful substrate for computations. Specifically, the model dissociates time and space and therefore can make use of Hebbian learning to learn spatiotemporal sequences. More general, the backbone as clustered organization might encode any variable *x* and enable downstream read-out neurons to learn and compute any function of this variable, *ϕ*(*x*).

## Methods

Excitatory neurons are modelled with the adaptive exponential integrate-and-fire model. A classical integrate-and-fire model is used for the inhibitory neurons. Excitatory to excitatory recurrent synapses are plastic under the voltage-based STDP rule [Clopath et al., 2010]. This enables the creation of neuronal clusters and a feedforward embedding. Normalization and weight bounds are used to introduce competition and keep the recurrent network stable. Synapses from inhibitory to excitatory neurons in the recurrent network are also plastic under a local plasticity rule [Vogels et al., 2011]. In general, it prevents runaway dynamics. The connections from the recurrent network to the read-out neurons are plastic under the same voltage-based STDP rule. However, potentiation of read-out synapses is linearly dependent on the strength of the synapses. There is no normalization here to allow a continuous weight distribution.

### Network dynamics

#### Recurrent network

A network with *N*^*E*^ excitatory (*E*) and *N*^*I*^ inhibitory (*I*) neurons is homogeneously recurrently connected with connection probability *p*. The weights are initialized such that the network is balanced.

#### Read-out neurons

The *N*^*E*^ excitatory neurons from the recurrent network are all-to-all connected to *N*^*R*^ excitatory read-out (*R*) neurons. This weight matrix is denoted by *W*^*RE*^. During learning, the read-out neurons receive supervisory input from *N*^*R*^ excitatory supervisor (*S*) neurons. The connection from supervisor neurons to read-out neurons is one-to-one and fixed, *w*^*RS*^. *N*^*R*^ interneurons (*H*) are one-to-one and recurrently connected to the read-out neurons with fixed connection strengths, *w*^*RH*^ and *w*^*HR*^ (see table 1 for the recurrent network and read-out parameters). The *E* to *E, I* to *E* and the *E* to *R* connections are plastic.

**Table 1:**
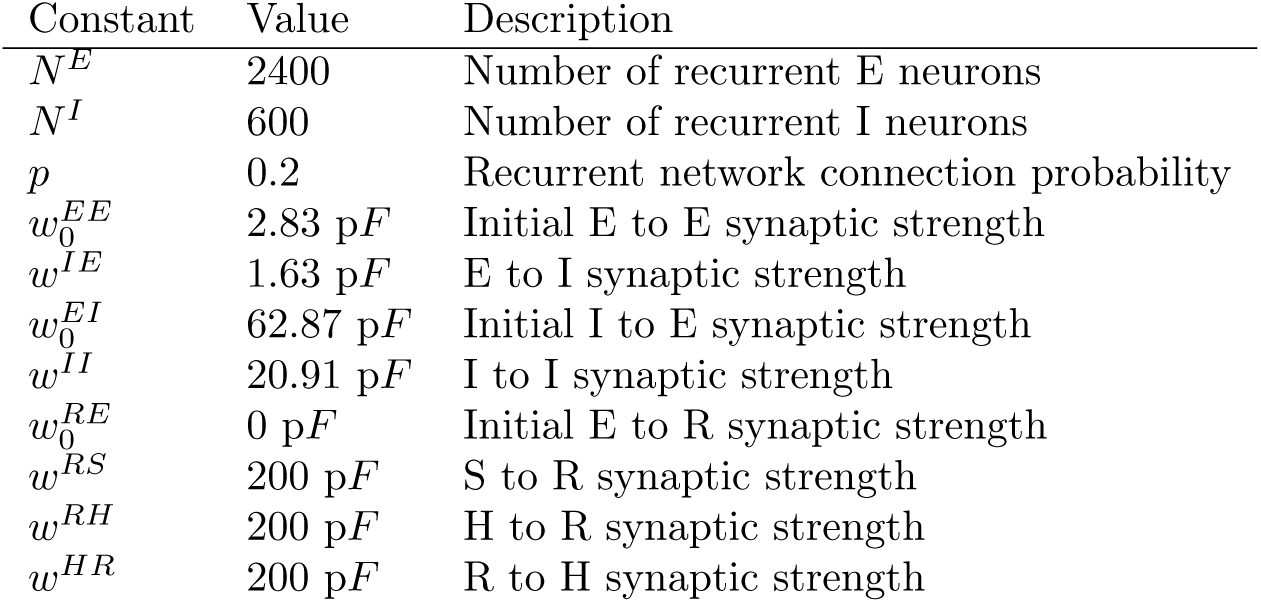
Initialization of network

#### Membrane potential dynamics

The membrane potential of the excitatory neurons (*V*^*E*^) in the recurrent network has the following dynamics:

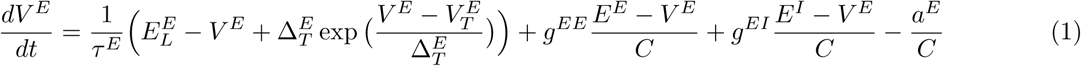

where *τ*^*E*^ is the membrane time constant, 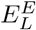 is the reversal potential, 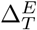 is the slope of the exponential, *C* is the capacitance, *g*^*EE*^, *g*^*EI*^ are synaptic input from excitatory and inhibitory neurons respectively and *E*^*E*^, *E*^*I*^ are the excitatory and inhibitory reversal potentials respectively. When the membrane potential diverges and exceeds 20 mV, the neuron fires a spike and the membrane potential is reset to *V*_*r*_. This reset potential is the same for all neurons in the model. There is an absolute refractory period of *τ*_*abs*_. The parameter 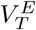 is adaptive for excitatory neurons and set to *V*_*T*_ + *A*_*T*_ after a spike, relaxing back to *V*_*T*_ with a time constant *τ*_*T*_:

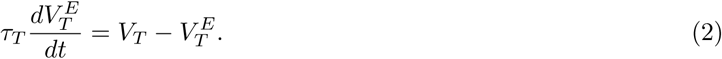

The adaptation current *a*^*E*^ for recurrent excitatory neurons follows:

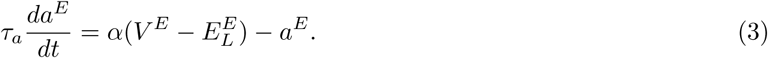

where *τ*_*a*_ is the time constant for the adaptation current and *α* is the slope. The adaptation current is increased with a constant *β* when the neuron spikes. The membrane potential of the read-out (*V* ^*R*^) neurons has no adaptation current:

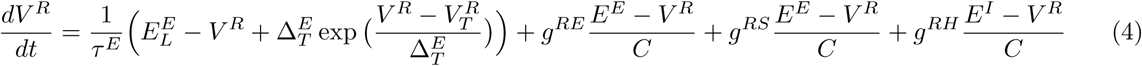

where 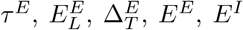 and *C* are as defined before. *g*^*RE*^ is the excitatory input from the recurrent network and supervisor neurons (supervisor input only non-zero during learning). *g*^*RS*^ is the excitatory input from the supervisor neuron (only non-zero during learning). *g*^*RH*^ is the inhibitory input from the interneuron (only non-zero during learning). The absolute refractory period is *τ*_*absR*_. The threshold 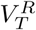 follows the same dynamics as 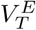, with the same parameters. The membrane potential of the supervisor neurons (*V*^*S*^) has no inhibitory input and no adaptation current:

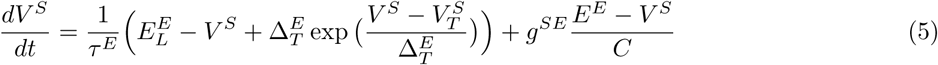

where the constant parameters are defined as before and *g*^*SE*^ is the external excitatory input from the target sequence. The absolute refreactory period is *τ*_*absS*_. The threshold 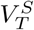 follows again the same dynamics as 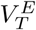, with the same parameters. The membrane potential of the inhibitory neurons (*V*^*I*^) in the recurrent network has the following dynamics:

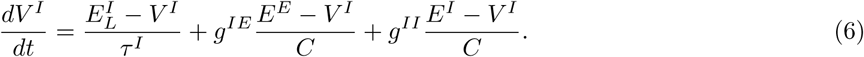

where *τ*^*I*^ is the inhibitory membrane time constant, 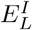 is the inhibitory reversal potential and *E*^*E*^, *E*^*I*^ are the excitatory and inhibitory resting potentials respectively. *g*^*EE*^ and *g*^*EI*^ are synaptic input from recurrent excitatory and inhibitory neurons respectively. Inhibitory neurons spike when the membrane potential crosses the threshold *V*_*T*_, which is non-adaptive. After this, there is an absolute refractory period of *τ*_*abs*_. There is no adaptation current. The membrane potential of the interneurons (*V*^*H*^) follow the same dynamics and has the same parameters, but there is no inhibitory input:

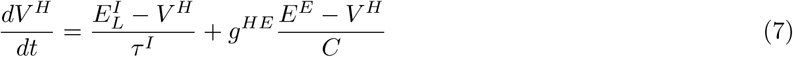

where the excitatory input *g*^*HE*^ comes from both the read-out neuron it is attached to and external input. After the threshold *V*_*T*_ is crossed, the interneuron spikes and an absolute refractory period of *τ*_*absH*_ follows. The interneurons inhibit the read-out neurons stronger when they receive strong inputs from the read-out neurons. This slows the potentiation of the read-out synapses down and keeps the synapses from potentiating exponentially (see table 2 for the parameters of the membrane dynamics).

**Table 2:** Neuronal membrane dynamics parameters

| Constant | Value | Description |
| --- | --- | --- |
| $\tau_E$ | 20 ms | E membrane potential time constant |
| $\tau_I$ | 20 ms | I membrane potential time constant |
| $\tau_{abs}$ | 5 ms | Refractory period of E and I neurons |
| $\tau_{absR}$ | 1 ms | R neurons refractory period |
| $\tau_{absS}$ | 1 ms | S neurons refractory period |
| $\tau_{absH}$ | 1 ms | H neurons refractory period |
| $E^E$ | 0 mV | excitatory reversal potential |
| $E^I$ | -75 mV | inhibitory reversal potential |
| $E_L^E$ | -70 mV | excitatory resting potential |
| $E_L^I$ | -62 mV | inhibitory resting potential |
| $V_r$ | -60 mV | Reset potential (for all neurons the same) |
| $C$ | 300 pF | Capacitance |
| $\Delta_T^E$ | 2 mV | Exponential slope |
| $\tau_T$ | 30 ms | Adaptive threshold time constant |
| $V_T$ | -52 mV | Membrane potential threshold |
| $A_T$ | 10 mV | Adaptive threshold increase constant |
| $\tau_a$ | 100 ms | Adaptation current time constant |
| $\alpha$ | 0 nS | Slope adaptation current of recurrent network neurons |
| $\beta$ | 1000 pA | Adaptation current increase constant of recurrent network neurons |

#### Synaptic dynamics

The synaptic conductance of a neuron *i* is time dependent, it is a convolution of a kernel with the total input to the neuron *i*:

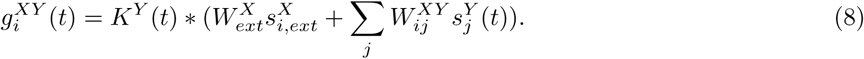

where *X* and *Y* denote two different neuron types in the model (*E, I, R, S* or *H*). *K* is the difference of exponentials kernel, 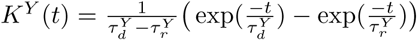, with a decay time *τ*_*d*_ and a rise time *τ*_*r*_ dependent only on whether the neuron is excitatory or inhibitory. There is no external inhibitory input to the supervisor and interneurons. During spontaneous activity, there is no external inhibitory input to the recurrent network and a fixed rate. The external input to the interneurons has a fixed rate during learning as well. The external input to the supervisor neurons is dependent on the specific learning task. There is no external input to the read-out neurons. The externally incoming spike trains 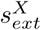 are generated from a Poisson process with rates 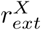. The externally generated spike trains enter the network through synapses 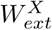 (see table 3 for the parameters of the synaptic dynamics).

**Table 3:** Synaptic dynamics parameters

| Constant | Value | Description |
| --- | --- | --- |
| $\tau_d^E$ | 6 ms | E decay time constant |
| $\tau_r^E$ | 1 ms | E rise time constant |
| $\tau_d^I$ | 2 ms | I rise time constant |
| $\tau_r^I$ | 0.5 ms | I rise time constant |
| $W_{ext}^E$ | 1.6 pF | External input synaptic strength to E neurons |
| $r_{ext}^E$ | 4.5 kHz | Rate of external input to E neurons |
| $W_{ext}^I$ | 1.52 pF | External input synaptic strength to I neurons |
| $r_{ext}^I$ | 2.25 kHz | Rate of external input to I neurons |
| $W_{ext}^S$ | 1.6 pF | External input synaptic strength to S neurons |
| $W_{ext}^H$ | 1.6 pF | External input synaptic strength to H neurons |
| $r_{ext}^H$ | 1.0 kHz | Rate of external input to H neurons |

### Plasticity

#### Excitatory plasticity

The voltage-based STDP rule is used [Clopath et al., 2010]. The synaptic weight from excitatory neuron *j* to excitatory neuron *i* is changed according to the following differential equation:

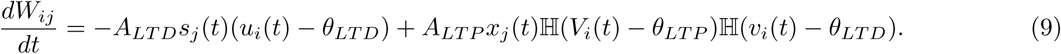

*A*_*LTD*_ and *A*_*LTP*_ are the amplitude of depression and potentiation respectively. *θ*_*LTD*_ and *θ*_*LTP*_ are the voltage thresholds to recruit depression and potentiation respectively, as ℍ (.) denotes the Heaviside function. *V*_*i*_ is the postsynaptic membrane potential, *u*_*i*_ and *v*_*i*_ are low-pass filtered versions of *V*_*i*_, with respectively time constants *τ*_*u*_ and *τ*_*v*_:

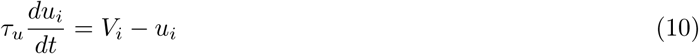

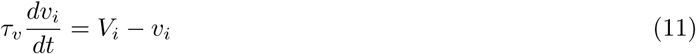

*s*_*j*_ is the presynaptic spike train and *x*_*j*_ is the low-pass filtered version of *s*_*j*_ with time constant *τ*_*x*_:

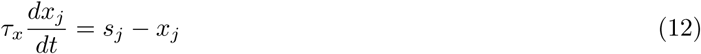

where the time constant *τ*_*x*_ is dependent on whether learning happens inside (*E* to *E*) or outside (*E* to *R*) the recurrent network. *s*_*j*_(*t*) = 1 if neuron *j* spikes at time *t* and zero otherwise. Competition between synapses is created in the recurrent network. This is done by a hard *L*1 normalization every 20 m*s*, keeping the sum of all weights onto a neuron constant: ∑ _*j*_*W*_*ij*_ = *K. E* to *E* weights have a lower and upper bound 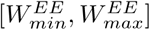. The minimum and maximum strengths are important parameters and determine the position of the dominant eigenvalues of *W*. Potentiation of the read-out synapses is weight dependent. Assuming that stronger synapses are harder to potentiate [Debanne et al., 1999], *A*_*LTP*_ reduces linearly with *W*^*RE*^:

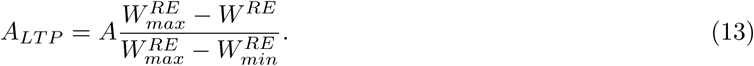

The maximum LTP amplitude *A* is reached when 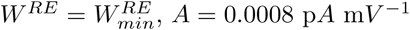, *A* = 0.0008 p*A* m*V* ^−1^ (see table 4 for the parameters of the excitatory plasticity rule).

**Table 4:** Excitatory plasticity parameters Description

| Constant | Value | Description |
| --- | --- | --- |
| $A_{LTD}$ | 0.0014 pA mV<sup>-2</sup> | LTD amplitude |
| $A_{LTP}$ | 0.0008 pA mV<sup>-1</sup> | LTP amplitude |
| $\theta_{LTD}$ | -70 mV | LTD threshold |
| $\theta_{LTP}$ | -49 mV | LTP threshold |
| $\tau_u$ | 10 ms | Time constant of low pass filtered membrane potential (LTD) |
| $\tau_v$ | 7 ms | Time constant of low pass filtered membrane potential (LTP) |
| $\tau_{xEE}$ | 3.5 ms | Time constant of low pass filtered spike train in recurrent network (LTP) |
| $\tau_{xRE}$ | 5 ms | Time constant of low pass filtered spike train for read-out synapses (LTP) |
| $W_{min}^{EE}$ | 1.45 pF | Minimum E to E weight |
| $W_{max}^{EE}$ | 32.68 pF | Maximum E to E weight |
| $W_{min}^{RE}$ | 0 pF | Minimum E to R weight |
| $W_{max}^{RE}$ | 25 pF | Maximum E to R weight |

#### Inhibitory plasticity

Inhibitory plasticity acts as a homeostatic mechanism, preventing runaway dynamics [Rhodes-Morrison et al., 2007, Vogels et al., 2011, Litwin-Kumar and Doiron, 2014, Zenke et al., 2015]. It is not necesseary for the stability of the sequential dynamics, but it is necessary for slow-switching dynamics (Fig S1). Excitatory neurons that fire with a higher frequency will receive more inhibition. The *I* to *E* weights are changed when the presynaptic inhibitory neuron or the postsynaptic excitatory neuron fires [Vogels et al., 2011]:

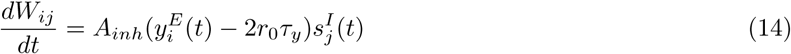

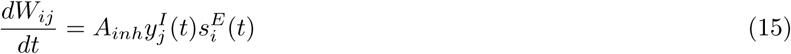

where *r*_0_ is a constant target rate for the postsynaptic excitatory neuron. *s*^*E*^ and *s*^*I*^ are the spike trains of the postsynaptic *E* and presynaptic *I* neuron respectively. The spike trains are low pass filtered with time constant *τ*_*y*_ to obtain *y*^*E*^ and *y*^*I*^ (as in equation 12). Table 5 shows parameter values for the inhibitory plasticity rule. The *I* to *E* synapses have a lower and upper bound 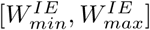.

**Table 5:**
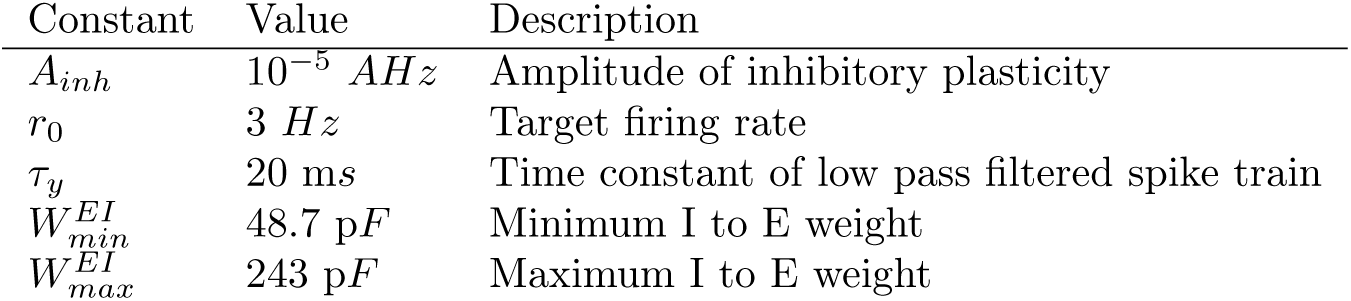
Inhibitory plasticity parameters

### Learning protocol

Learning happens in two stages. First a feedforward structure is embedded in the recurrent network that produces reliable sequential dynamics. Once this dynamics is established connections to read-out neurons can be learned. Read-out neurons are not interconnected and can learn in parallel.

#### Recurrent network

The network is divided in disjoint clusters of 80 neurons. The clusters are sequentially stimulated for a time duration of 60 minutes by a large external current where externally incoming spikes are drawn from a Poisson process with rate 18 k*Hz*. This high input rate does not originate from a single external neuron but rather assumes a large external input population. Each cluster is stimulated for 10 m*s* and in between cluster stimulations there are 5 m*s* gaps. During excitatory stimulation of a cluster, all other clusters receive an external inhibitory input with rate 4.5 k*Hz* and external input weight 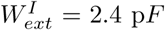. There is a periodic boundary condition. The last cluster activates the first cluster again. After the sequential stimulation, the network is spontaneously active for 60 minutes. The connectivity stabilizes during the spontaneous dynamics. The recurrent weight matrix of the large network (Fig 5) is learned using the same protocol. The recurrent weight matrix reaches a stable structure after three hours of sequential stimulation followed by three hours of spontaneous dynamics. Parameters that change are summarized in table 6. For slow-switching dynamics, a similar protocol is followed, with minor changes (Fig S1). The weight matrix used to plot the spectrum of the recurrent network in Figs 2 and S2 is 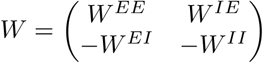.

**Table 6:** Large recurrent network changed parameters

| Constant | Value | Description |
| --- | --- | --- |
| $N^E$ | 6400 | Number of recurrent E neurons |
| $N^I$ | 1600 | Number of recurrent I neurons |
| $w_0^{EE}$ | 1.73 pF | baseline E to E synaptic strength |
| $w^{IE}$ | 1.20 pF | E to I synaptic strength |
| $w_0^{EI}$ | 38.50 pF | Initial I to E synaptic strength |
| $w^{II}$ | 12.80 pF | I to I synaptic strength |
| $W_{min}^{EE}$ | 1.27 pF | Minimum E to E weight |
| $W_{max}^{EE}$ | 30.5 pF | Maximum E to E weight |
| $W_{min}^{EI}$ | 40 pF | Minimum I to E weight |
| $W_{max}^{EI}$ | 200 pF | Maximum I to E weight |
| $W_{max}^{RE}$ | 15 pF | Maximum E to R weight |

#### Read-out network

During learning of the read-out synapses, external input drives the supervisor and interneurons. The rate of the external Poisson input to the supervisor neurons reflects the sequence that has to be learned. The rate is normalized between 0 k*Hz* and 10 k*Hz*. During learning, *W*^*RE*^ changes. After learning, the external input to the supervisor and interneurons is turned off and both stop firing. The read-out neurons are now solely driven by the recurrent network. Plasticity is frozen in the read-out synapses after learning. With plasticity on during spontaneous dynamics, the read-out synapses would continue to potentiate because of the coactivation of clusters in the recurrent network and read-out neurons. This would lead to read-out synapses that are all saturated at the upper weight bound.

#### Simulations

The code used for the training and simulation of the recurrent network is build on top of the code from [Litwin-Kumar and Doiron, 2014] in Julia. The code used for learning spatiotemporal sequences using read-out neurons is written in Matlab. Recurrent networks learned in Julia can be saved and loaded into the Matlab code, or networks with a manually set connectivity can be explored as in [Schaub et al., 2015]. Forward Euler discretization with a time step of 0.1 m*s* is used. The code will be made public on ModelDB after publication.

## Supplementary material

### Linear rate model: analysis of the spectrum

A linear rate model can give insight into the dynamics of a large nonlinear structured spiking network [Schaub et al., 2015]. Sequential dynamics in a linear rate network can emerge from the same feedforward structure as studied in this paper. The dynamics of a simplified rate model follows:

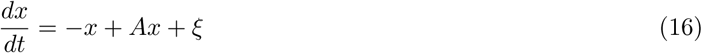

where *x* is a multidimensional variable consisting of the rates of all excitatory and inhibitory groups, *A* is the weight matrix and *ξ* is noise. A minimal model for sequential firing consists of three groups of excitatory neurons and one group of inhibitory neurons (Fig S3). We are interested in a general connectivity structure:

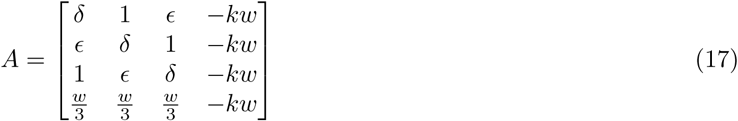

where *w* = *δ* + *ϵ* + 1, *ϵ* > 1 for sequential dynamics and *k* > 1 for balance. The Schur decomposition *A* = *UTU*^*T*^ gives both the eigenvalues and an orthonormal basis for the eigenvectors:

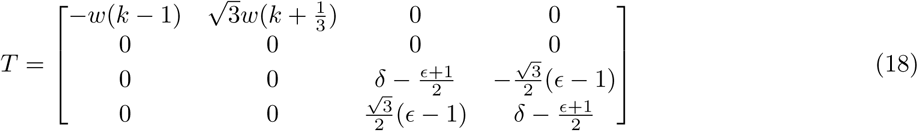

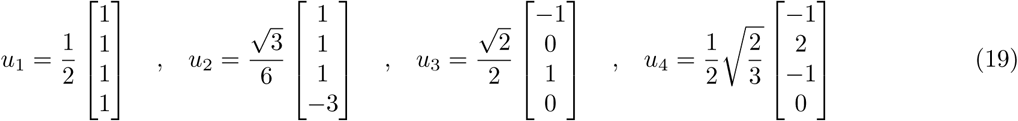

where the basis for the complex conjugate eigenvalues is a plane and can be rotated (for *ϵ* > 1). The first mode, *u*_1_, decays fast and uniformly over the different neuronal groups. The second mode, *u*_2_, decays slower and indicates the interplay between excitatory and inhibitory groups. Mode three and four span the eigenspace for the complex conjugate pair, {*u*_3_, *u*_4_}. This space is localized on the three excitatory groups. It shows that an increase of activity in one group is coupled with decreased activities in other groups. The real part 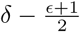 should be smaller than one for linear stability and closer to one means a slower decay of this mode. Importantly, the imaginary part is linearly dependent on the amount of feedforward structure, *ϵ* (Fig S3A). This leads to an oscillation and essentially determines the frequency of sequential switching.

**Figure S1:**
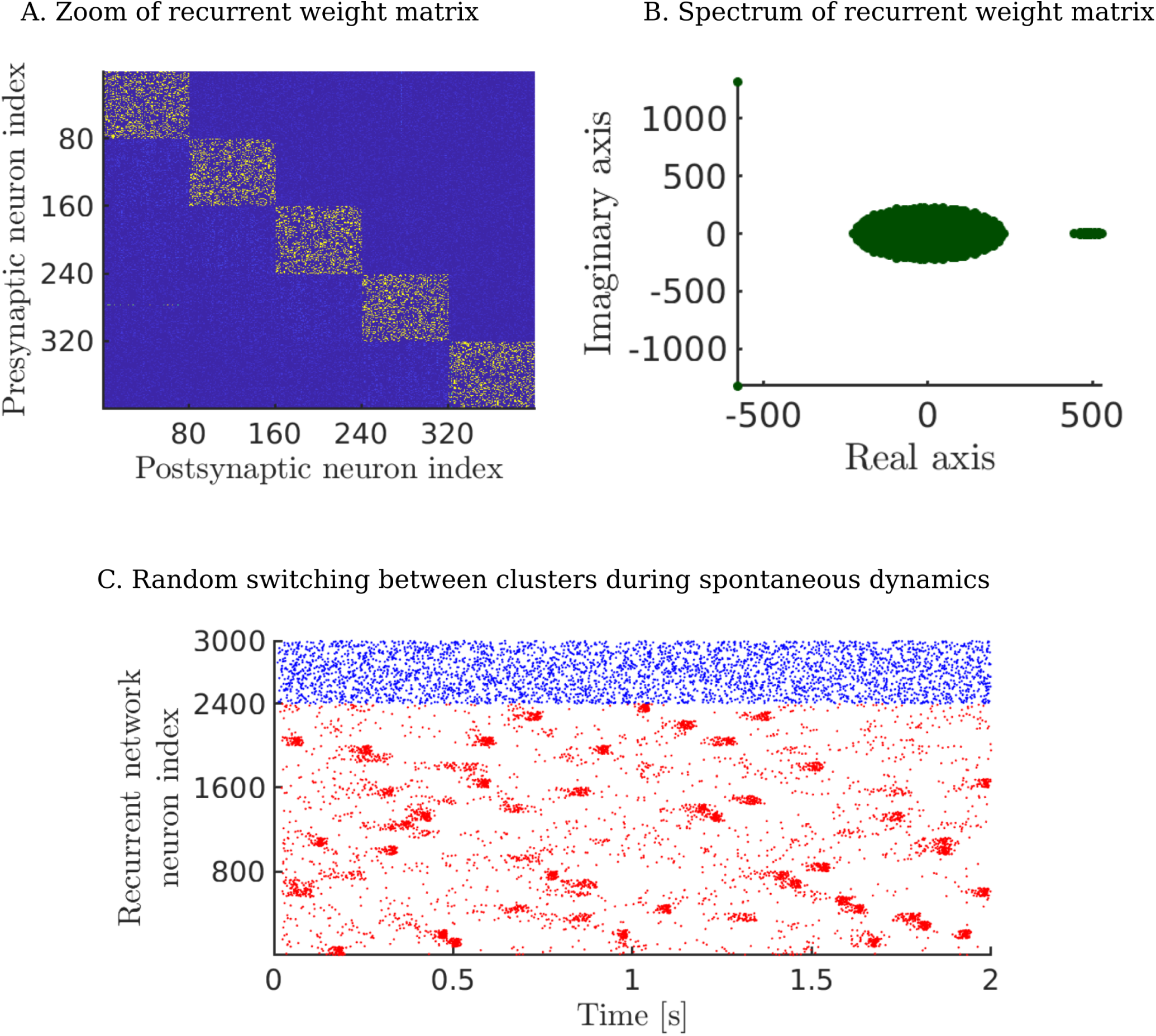
Slow-switching dynamics. The recurrent network is stimulated with external input that is spatially clustered, but temporally uncorrelated. Each cluster is stimulated for 50 m*s*, with 50 m*s* gaps in between stimulations. The rate of external stimulation is 18 k*Hz* during training and there is no simultaneous inhibition of other clusters. This is repeated for 20 minutes after which the network stabilizes during 20 minutes of spontaneous activity. (A.) A diagonal structure is embedded in the recurrent network. Since there are no temporal correlations in the external input, there is no off-diagonal structure. (B.) The spectrum shows an eigenvalue gap. This indicates the emergence of a slower time scale. The leading eigenvalues do not have an imaginary part, pointing at the absence of feedforward structure and thus there is no sequential dynamics. (C.) Under a regime of spontaneous dynamics (i.e. uncorrelated Poisson inputs), the clusters are randomly reactivated. The high adaptation means a neuron fires only once during each reactivation.

**Figure S2:**
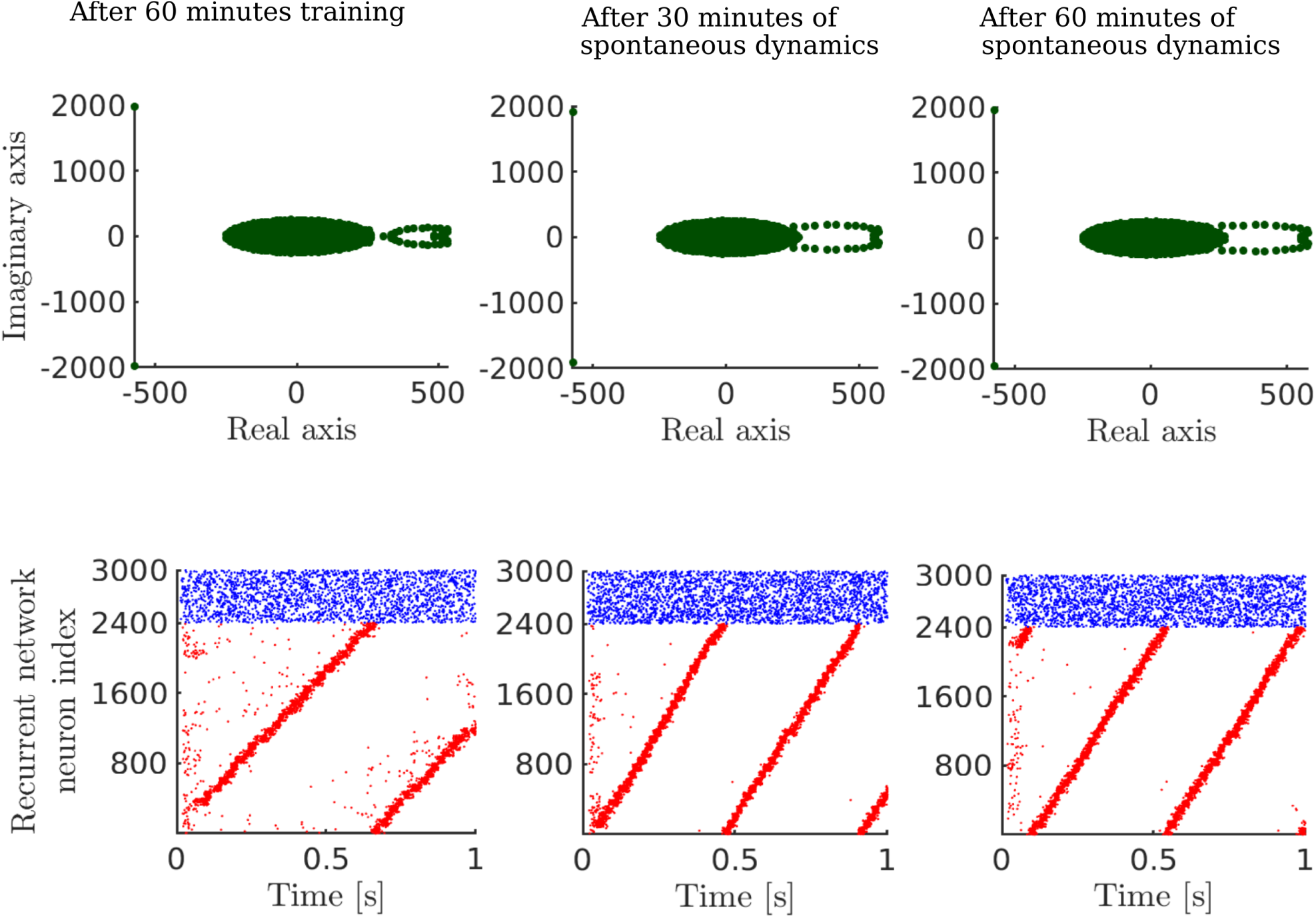
The feedforward embedding is stable. After 60 minutes of training (i.e. sequential stimulation as described in the methods), the network stabilizes during spontaneous activity. During the first 30 minutes of spontaneous dynamics, the connectivity still changes. More specifically, the imaginary parts of the leading eigenvalues increase. This leads to a higher switching frequency and as such a smaller period in the sequential activity. After around 30 minutes, a fixed point is reached. The first row shows the spectra of the weight matrix at different times. The second row shows the spike trains at those times, for one second of spontaneous activity.

**Figure S3:**
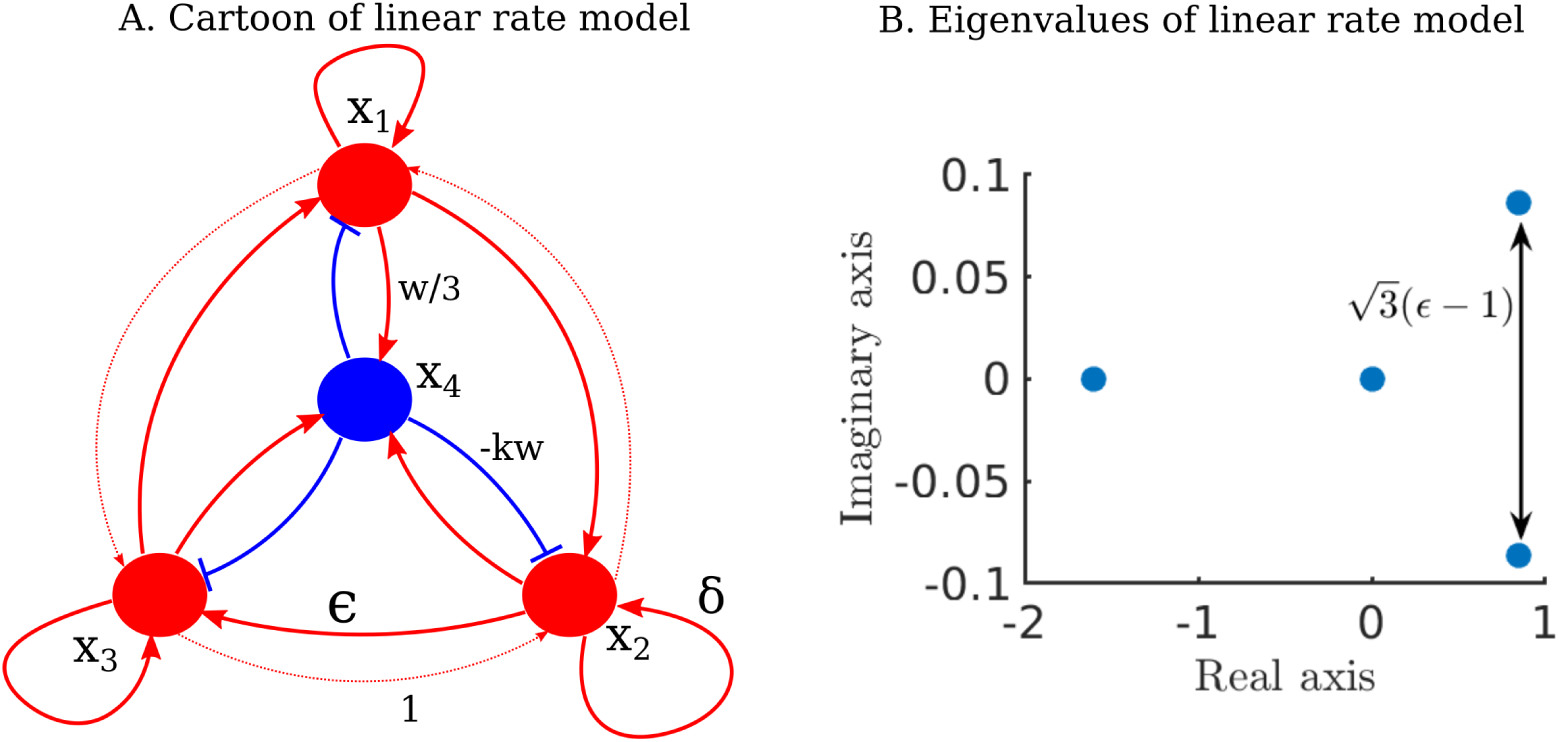
(A.) Cartoon of simple linear model, the cyclic connections are stronger clockwise than anticlockwise. (B.) Eigenvalue spectrum shows a conjugate complex eigenvalue pair. The imaginary part scales linearly with *E*.

**Figure S4:**
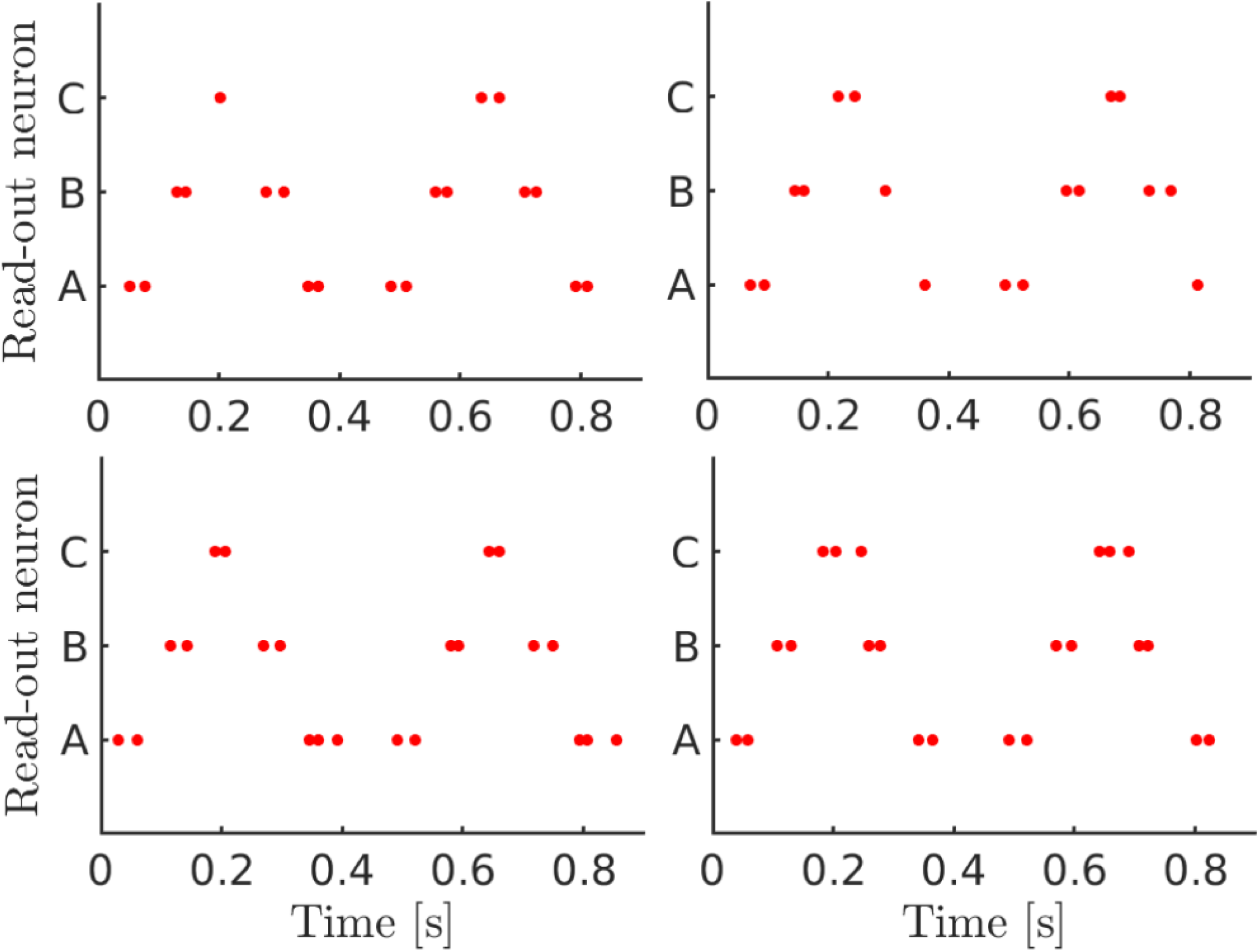
The sequence *ABCBA* is relearned four times for 12 seconds each. Before relearning, the read-out weight matrix *W*^*RE*^ was always reset. When active, read-out neurons fire two spikes on average +*/*− one spike. This variability is a consequence of the noisy learning process.

**Figure S5:**
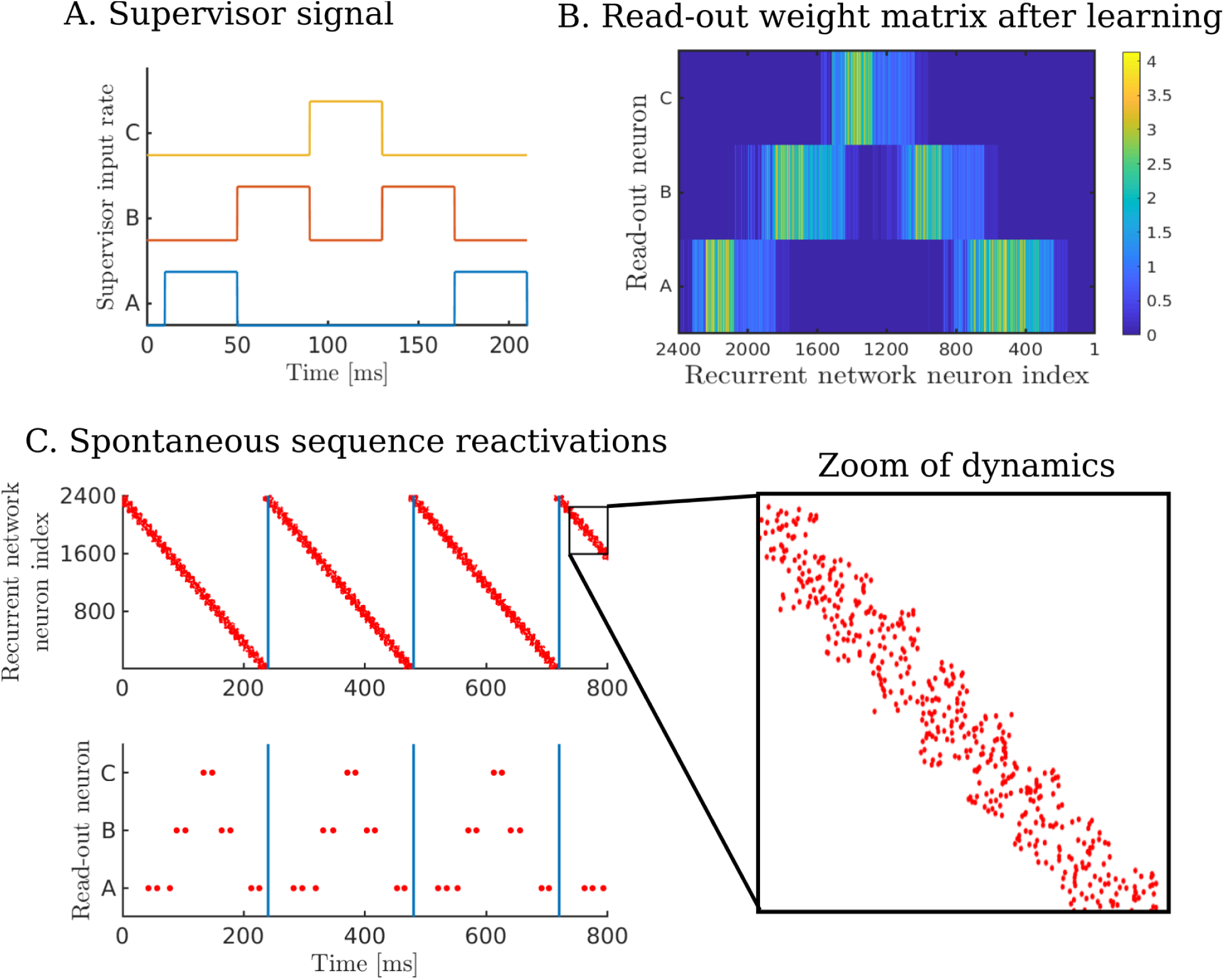
The dynamics of the recurrent network is forced to overlap. 2400 excitatory neurons are grouped in 30 clusters of 80 neurons. Whereas in the trained networks each cluster is discretely active, here their activity overlaps. Neurons in cluster *i* can fire in time bin [*t*_*i-*1_, *t*_*i*_] and in time bin [*t*_*i*_, *t*_*i*+1_]. Learning with such overlapping dynamics is still possible. (A.) The non-markovian sequence *ABCBA* is learned where each letter corresponds to a 40 m*s* stimulation of the corresponding supervisor neuron. (B.) The read-out weight matrix after learning for 10 seconds. (C.) The forced dynamics in the recurrent network drives spiking activity in the read-out neurons. The zoom of activity in the recurrent network shows the overlapping dynamics.

